# Co-evolution of Eukaryotic-like Vps4 and ESCRT-III Subunits in the Asgard Archaea

**DOI:** 10.1101/2020.05.05.080093

**Authors:** Zhongyi Lu, Ting Fu, Tianyi Li, Yang Liu, Siyu Zhang, Jinquan Li, Junbiao Dai, Eugene V Koonin, Guohui Li, Huiying Chu, Meng Li

## Abstract

The emergence of the endomembrane system is a key step in the evolution of cellular complexity during eukaryogenesis. The Endosomal Sorting Complex Required for Transport (ESCRT) machinery is essential and required for the endomembrane system functions in eukaryotic cells. Recently, genes encoding eukaryote-like ESCRT protein components have been identified in the genomes of Asgard archaea, a newly proposed archaeal superphylum that is thought to include the closest extant prokaryotic relatives of eukaryotes. However, structural and functional features of Asgard ESCRT remain uncharacterized. Here we show that Vps4, Vps2/24/46, and Vps20/32/60, the core functional components of the Asgard ESCRT, co-evolved eukaryote-like structural and functional features. Phylogenetic analysis shows that Asgard Vps4, Vps2/24/46, and Vps20/32/60 are closely related to their eukaryotic counterparts. Molecular dynamic simulation and biochemical assays indicate that Asgard Vps4 contains a eukaryote-like Microtubule Interacting and Transport (MIT) domain that binds the distinct type-1 MIT Interacting Motif and type-2 MIT Interacting Motif in Vps2/24/46, and Vps20/32/60, respectively. The Asgard Vps4 partly, but much more efficiently than homologs from other archaea, complements the *vps4* null mutant of *Saccharomyces cerevisiae*, further supporting the functional similarity between the membrane remodeling machineries of Asgard archaea and eukaryotes. Thus, this work provides evidence that the ESCRT complexes from Asgard archaea and eukaryotes are evolutionarily related and functionally similar. Thus, despite the apparent absence of endomembranes in Asgard archaea, the eukaryotic ESCRT seems to have been directly inherited from an Asgard ancestor, to become a key component of the emerging endomembrane system.

**IMPORTANCE:** The discovery of Asgard archaea has changed the exiting ideas on the origins of Eukaryotes. Researchers propose that eukaryotic cells evolve from Asgard archaea. This hypothesis partly stems from the presence of multiple eukaryotic signature proteins in Asgard archaea, including homologues of ESCRT proteins that are essential components of the endomembrane system in eukaryotes. However, structural and functional features of Asgard ESCRT remain unknown. Our study provides evidence that Asgard ESCRT is functionally comparable to the eukaryotic counterparts suggesting that, despite the apparent absence of endomembranes in archaea, eukaryotic ESCRT was inherited from an Asgard archaeal ancestor, alongside the emergence of endomembrane system during eukaryogenesis.

## INTRODUCTION

Eukaryogenesis is a major, long-standing puzzle in evolutionary biology because the specifics of the evolutionary process leading to the eukaryotic cellular complexity are far from being clear. One of the key distinctions of eukaryotic cells from the cells of prokaryotes is the presence in the former of the sophisticated endomembrane system. Undoubtedly, the emergence of the endomembrane system was a milestone event in eukaryogenesis because it is a pre-requisite of the intracellular compartmentalization which is a hallmark of eukaryotic cells (1). The Endosomal Sorting Complex Required for Transport (ESCRT) machinery is an essential component of the eukaryotic endomembrane system that, as such, has been thought to be restricted to eukaryotic cells (2). For instance, *Saccharomyces cerevisiae* ESCRT consists of five main subcomplexes: ESCRT-0, -I, -II, -III, and Vps4 (3-5). Of these, the Vps4 and ESCRT-III subunits are central players in ESCRT function that mediates remodelling and scission of endomembranes (6, 7). The ESCRT-III subunits can be further divided into two classes, termed Vps2/24/46 and Vps20/32/60, and both participate in either directly or indirectly forming membrane-bound polymeric assemblies that sever membrane necks (8). On the other hand, Vps4, an ATPase, promotes ATP-dependent disassembly of the ESCRT-III polymers, thus ensuring the ESCRT-III subunits turnover. Several studies have shown that the N-terminal Microtubule Interacting and Transport (MIT) domain of Vps4 recognizes and interacts with the type-1 MIT Interacting Motif (MIM1) that is present in the Vps2/24/46 class ESCRT-III subunits and type-2 MIT Interacting Motif (MIM2) present in the Vps20/32/60 subunits. These recognition models are essential for the biological function of ESCRT-III and Vps4 (9-11).

The cell division (Cdv) systems discovered in some archaeal orders, such as Sulfolobales and Desulfurococcales within the TACK (Thaumarchaeota, Aigarchaeota, Crenarchaeota, and Korarchaeota) superphylum include a homolog of eukaryotic Vps4 (CdvC) and several homologs of eukaryotic ESCRT-III subunits (CdvBs) (12, 13). Given these homologies and because in eukaryotes, the MIT-MIM2 interactions occurred between CdvC and CdvB (12, 14, 15), the creanarchaeal Cdv system has been proposed to be the evolutionary ancestor of eukaryotic ESCRT (16). However, this evolutionary relationship remains uncertain. One reason for the uncertainty is that CdvBs lack the well-characterized MIM1, and the absence of the MIT-MIM1 interaction is likely to reflect major functional differences between crenarchaeal Cdv and eukaryotic ESCRT (12, 17). Such differences might indicate that, although the two systems consist of homologous subunits, the Cdv system is not the direct ancestor of eukaryotic ESCRT.

The recently discovered Asgard archaea (including Lokiarchaeota, Thorarchaeota, Heimdallarchaeota, Odinarchaeota, and Helarchaeota) have been proposed to include the closest archaeal relatives of eukaryotes. This proposition stems, partly, from the findings that the Asgard genomes encode a broad repertoire of Eukaryotic Signature Proteins (ESPs) that are far more prevalent in Asgard than they are in other archaea (18-21). Among these ESPs are highly conserved homologs of eukaryotic ESCRT-I, -II, -III, and Vps4. Notably, the presence of these proteins in Asgard archaea that was originally demonstrated on metagenomics assemblies has been confirmed by analysis of the first closed Asgard genome, ruling out the possibility of a eukaryotic contamination (12, 18, 19, 22).

Here, we explore the phylogenetic relationships among the ESCRT-III components, reconstitute and biochemically characterize the Asgard Vps4, and test its potential biological function in the heterologous *S. cerevisiae* endomembrane system. The combined phylogenetic, genetic and biochemical analyses reveal close relationships between the ESCRT-III subunits and Vps4 of Asgard archaea and eukaryotes, to the exclusion of other archaea.

## RESULTS

### Eukaryotic-like ESCRT-III subunits and Vps4 in Asgard archaea

Given that the ESCRT-III subunits are tightly linked to the functional complexity of ESCRT (12), we first performed a detailed sequence comparison and phylogenetic analysis of the Vps2/24/46 and Vps20/32/60 as well as the Vps4 ATPase from Asgard archaea based on the available genomic data(18, 19). In the unrooted maximum-likelihood phylogenetic tree Vps2/24/46 and Vps20/32/60, the Asgard proteins form a cluster with eukaryotic homologs that is separated from the archaeal (TACK) CdvB cluster by a long branch (Fig. 1A, Fig. S1 and Table S1), supporting the notion that Asgard archaea possess “eukaryote-like” ESCRT-III subunits. All the Asgard Vps20/32/60 proteins form a strongly supported clade with the eukaryotic Vps20/32/60 which is compatible with a direct ancestral relationship. The Asgard Vps2/24/46 proteins formed three clades one of which (Odinarchaeota, Lokiarchaeota, and Thorarchaeota) clustered with the eukaryotic homologs whereas the remaining two (Heimdallarchaeota) placed near the root of the Asgard-eukaryote branch (Fig. 1A). This tree topology is likely to result from acceleration of evolution in Heimdallarchaeota.

**FIG 1.**
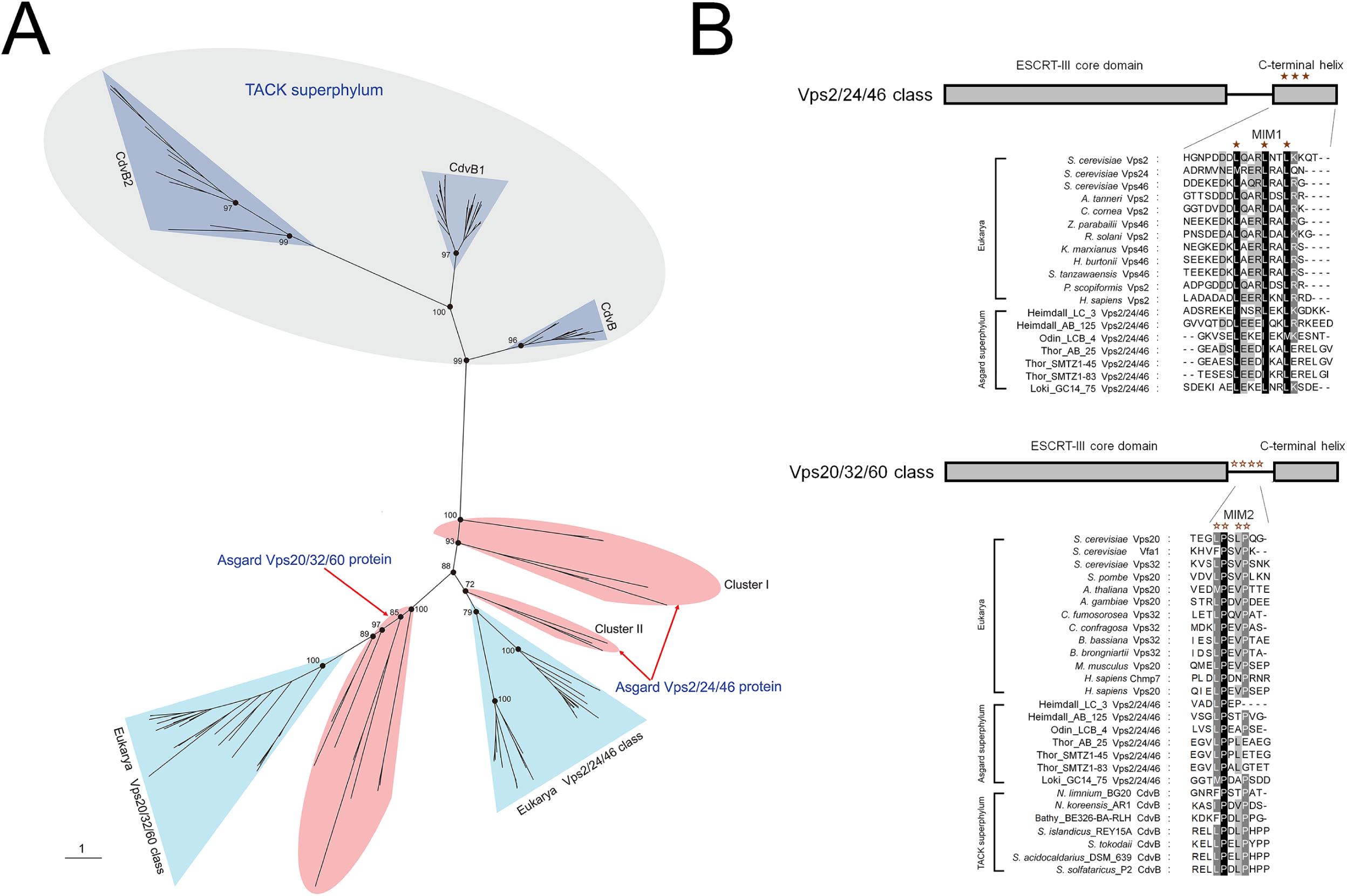
Phylogenetic and amino acid sequence analysis of the ESCRT- III-related subunits in archaea and eukarya. (A) Unrooted maximum- likelihood phylogenetic tree of the ESCRT-III-related subunits in archaea and eukarya. The information of the Asgard Vps2/24/46 and Vps20/32/60 can be found in Table S1. Part of the bootstrap values are shown on nodes. (B) Predicted MIM1 and MIM2 in Asgard Vps2/24/46 and Vps20/32/60, respectively. The information of proteins used here can be found in Table S1. The ESCRT-III core domain, C-terminal helix, and MIM1 and MIM2 are presented.

In addition to the phylogenetic results, we found that the Asgard Vps2/24/46 contained leucine-rich motifs located in the C-terminal helix and resembling the C-terminal MIM1 that are conserved in eukaryotes although some leucine residues were substituted by isoleucine in the Asgard homologs (Fig. 1B) (9). The C-terminal regions of the Asgard Vps20/32/60 contain proline-rich motifs that resemble MIM2 although do not fully conform to the MIM2 consensus in eukaryotes and TACK archaea (10, 11). Taken together, the results of phylogenetic analysis and motif search for ESCRT-III subunits not only demonstrate the Asgard-eukaryote affinity but also show that the ancestors of the Vps2/24/46 and Vps20/32/60 groups have already diverged in Asgard archaea, antedating eukaryogenesis.

It appears likely that Vps4 structurally and functionally co-evolved with ESCRT-III subunits in Asgard archaea. To explore the evolution of Vps4, an unrooted maximum-likelihood phylogenetic tree was constructed for the group of ATPases including CdvC from the TACK superphylum, Asgard Vps4, and the so called eukaryotic “meiotic clade” comprised of Vps4, Katanin 60, and Spastin (23). As in the ESCRT-III subunit tree, the Asgard Vps4 formed a branch with the eukaryotic homolog that was separated by a long, strongly supported branch from the archaeal CdvC branch (Fig. 2A and Fig. S2). The Asgard Vps4 did not form a single clade, but rather four clades, all of which were located close to the root of the Asgard-eukaryote branch.

**FIG 2.**
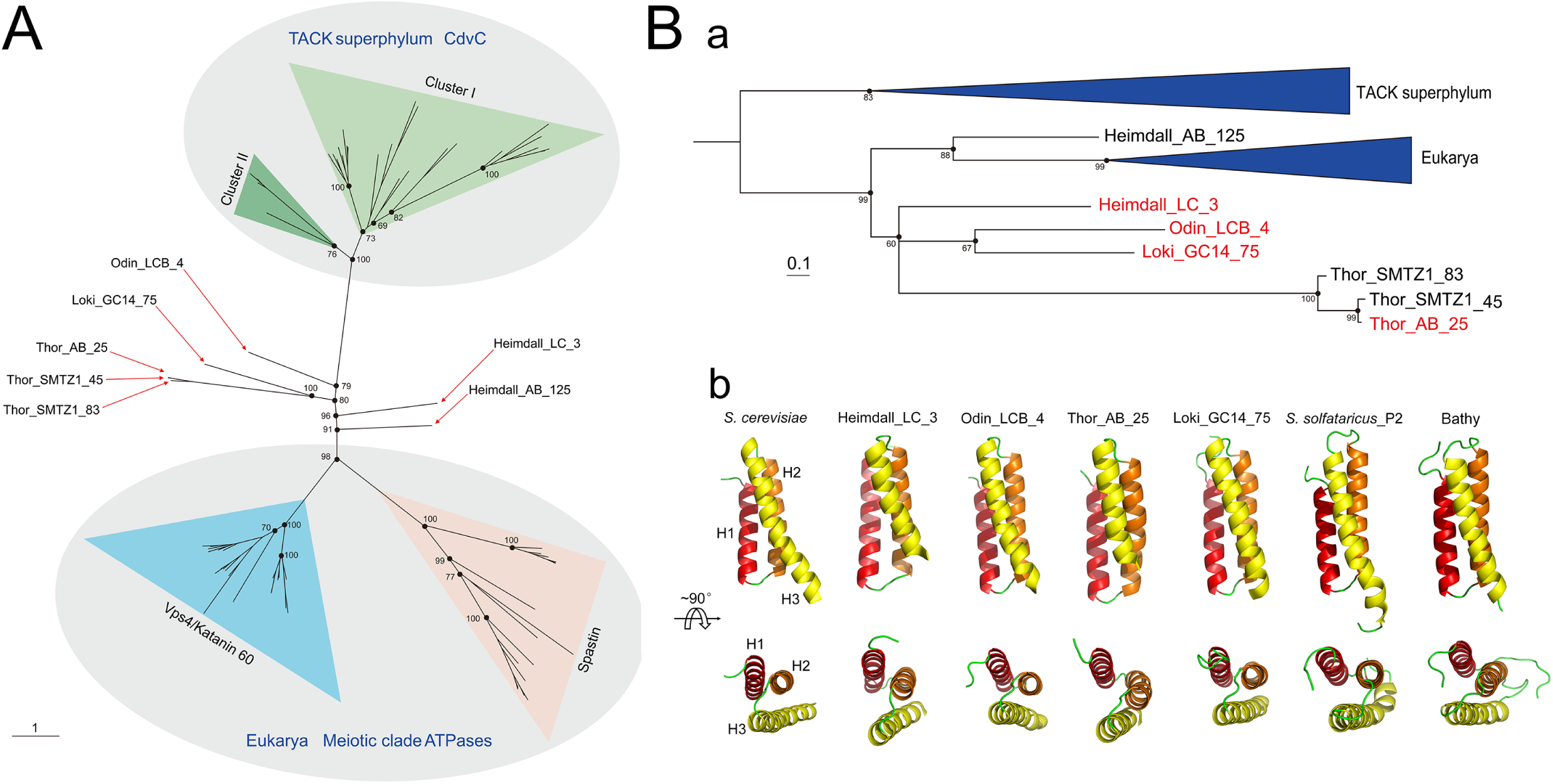
Phylogenetic and structural analysis of the Asgard Vps4. (A) Unrooted maximum likelihood phylogenetic analysis of the Vps4- related in archaea and eukarya. The information of the Asgard Vps4 can be found in Table S1. Part of the bootstrap values are shown on nodes. (B) Phylogenetic (a) and structural (b) analysis of the Asgard Vps4 MIT domain. The sequences of CdvC MIT domain are used as the outgroup to further confirm the phylogenetic relationship of the MIT domain in eukaryotic and Asgard Vps4. The antiparallel three-helix bundle of MIT domains is shown explicitly.

Despite their high divergence demonstrated by the lack of monophyly in the phylogenetic tree (Fig. 2A and Fig. S2), all Asgard Vps4 contain the eukaryotic-like “arginine collar” that consists of three conserved arginine residues (Fig. S3A). In eukaryotes, this motif is located in the pore loop 2 of Vps4 and is involved in the ESCRT-III filaments translocation to the central pore of the Vps4 hexamer for disassembly (Fig. S3B) (24, 25).

Because Vps4 recognizes ESCRT-III subunits via the MIT domain, we specifically analyzed the phylogeny of the MIT domains of the Vps4 proteins from Asgard archaea, eukaryotes, and TACK archaea. The tree demonstrates a clear affiliation of Asgard with eukaryotes that, in this case, form a clade with one of the MIT domain from Heimdallarchaeota (Fig. 2B and Fig. S4). Affiliation with Heimdallarchaeota has been previously observed for many Asgard genes(19, 26, 27).

To further structurally characterize MIT domain in Asgard Vps4, we constructed stable models of full length Vps4 from Heimdallarchaeota_LC_3, Odinarchaeota_LCB_4, Thorarchaeota_AB_25, and Lokiarchaeum_GC14_75 using homology modeling and molecular dynamics simulation, and compared these with the *S. cerevisiae* Vps4 structure. As a control for the Asgard Vps4, we include CdvC from *Sulfolobus solfataricus*_P2 (cluster I in Fig. 2A) and Bathyarchaeota (cluster II in Fig. 2A). All MIT domains of Asgard Vps4 and TACK CdvC adopted a three-helix bundle structure that is closely similar to the *S. cerevisiae* MIT domain structure although the helices in both the Asgard and TACK structures are somewhat shorter than in the *S. cerevisiae* structure (Fig. 2Bb and Fig. S5).

Taken together, the above data suggest that the evolution of Asgard Vps4, especially their MIT domain, was accompanied by the functional divergence of the ESCRT-III subunits. Thus, although the Asgard Vps4 proteins are highly diverged, the results of sequence comparison, phylogenetic analysis and structural modeling are compatible with coevolution of Vps4 with ESCRT-III subunits and an ancestral relationship between the membrane remodeling machineries of Asgard and eukaryotes. Furthermore, it can be predicted that Asgard Vps2/24/46 and Vps20/32/60 form ESCRT-III-like filaments similar to those in eukaryotes.

### Interactions between Asgard Vps4 and ESCRT-III subunits

As previously described, unlike the CdvBs, Asgard Vps2/24/46 and Vps20/32/60 share the same eukaryotic ESCRT-III secondary structure, and these organizations are probably responsible for their ability bind to Vps4 like their eukaryotic counterparts(12). The isothermal titration calorimetry (ITC) was adopted to verify that the MIT domain of Asgard Vps4 bind to Vps2/24/46 and Vps20/32/60, respectively (Fig. S6). To characterize the interactions between Asgard Vps4 and ESCRT-III subunits, the respective binding free energies were estimated by MM-GBSA calculations (Table S2A) (28, 29). The binding free energies of Vps4-Vps2/24/46 in Heimdall_LC_3, Odin_LCB_4, Thor_AB_25, and Loki_GC14_75 were calculated as -39.02, -61.85, -71.81, -72.24 kcal/mol, respectively. All these values, although compatible with stable binding, are lower than the binding free energy of Vps4-Vps2 (−82.98 kcal/mol) in *S. cerevisiae*, suggesting that the affinity of Asgard Vps4 for Vps2/24/46 is weaker than that of *S. cerevisiae* Vps4 for Vps2. The binding free energies for Asgard Vps4- Vps20/32/60 differed to a greater degree indicating variation in the affinities (Table S2B). The Thor_AB_25 value of -119.97 kcal/mol was substantially greater than the binding free energy of the Vps4-Vps20 interaction in *S. cerevisiae* (−88.30 kcal/mol), the values for Heimdall_LC_3 (−81.43 kcal/mol) and Loki_GC14_75 (−88.89 kcal/mol) comparable to those for *S. cerevisiae*, and that for Odin_LCB_4 (−49.53 kcal/mol) much lower than in *S. cerevisiae*.

We further analyzed the structural basis for the MIT domain of Asgard Vps4 binding to the putative MIM1 and MIM2 of Vps2/24/46 and Vps20/32/60, respectively, by using MM-GBSA calculations(28). The key amino acids that contribute to the Vps4 MIT domain binding to the Vps2/24/46 MIM1 in Heimdall_LC_3, Odin_LCB_4, and Thor_AB_25 are mainly located in Helix 2 and Helix 3 of the MIT domain similar to the location of MIM1-interacting residues in *S. cerevisiae* Vps4 (Fig. 3A and Table S3). These findings are consistent the MIM1 peptide biding at the interface between Helix 2 and Helix 3 of the MIT domain as observed in eukaryotes (10, 30). In Loki_GC14_75, the key amino acid residues are located in Helix 1 and Helix 2, suggesting a distinct interaction mode.

**FIG 3.**
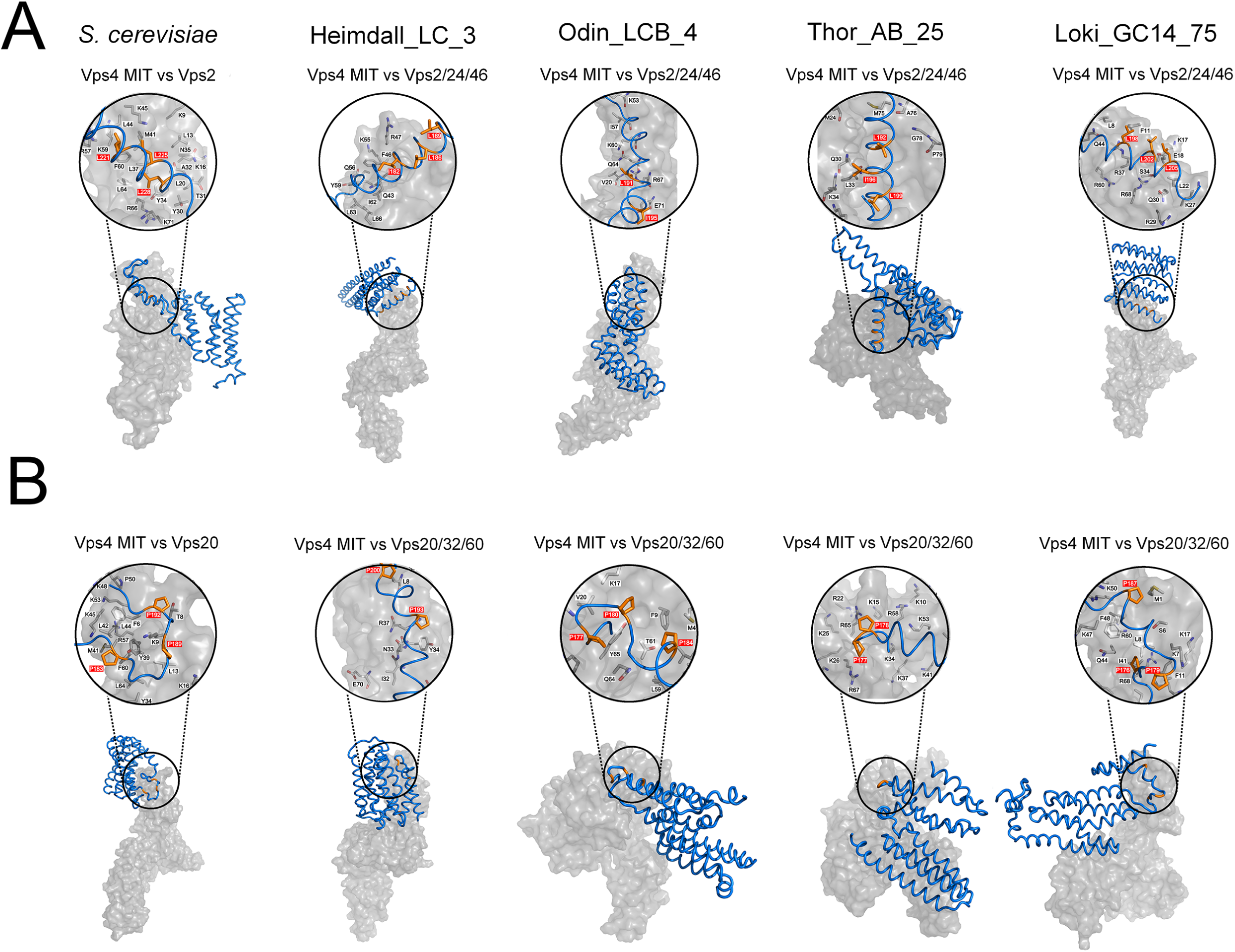
Comparison of the Vps4 (surface representation, grey) in complex with ESCRT-III subunits (ribbon representation, blue) in Asgard archaea. The MIM1 and MIM2 are shown in orange (stick representation, orange) and highlighted in red in close-up views (space filling representation). The black letters indicated main residues in MIT domains that contribute to the interaction. The Vps4 MIT domain in complex with (A) Vps2/24/46 and (B) Vps20/32/60 subunits in *S. cerevisiae*, Heimdall_LC_3, Odin_LCB_4, Thor_AB_25, and Loki_GC14_75 are indicated.

The key residues involved in the MIT-Vps20/32/60 interactions are spread among Helix 1, Helix 2, and Helix 3, in positions closely similar to those involved in the MIT-Vps20 interactions in *S. cerevisiae* (Fig. 3B and Table S4). Thus, the MIM2 peptide is predicted to bind the grooves formed by the three-helix bundle rather than Helix 1 and Helix 3 only as also observed for the eukaryotic ESCRT-III (10, 11). Taken together, these findings indicate that the interactions of the Asgard Vps4 MIT domain with the MIM1 (in Vps2/24/46) and MIM2 (in Vps20/32/60) motifs closely resemble the corresponding interactions in eukaryotes.

### Asgard Vps4 phenotypically complements *vps4* null mutation in *S. cerevisiae*

We further sought to determine whether the Asgard and eukaryotic Vps4 ATPases were functionally interchangeable. To this end, Heimdall_LC_3, Odin_LCB_4, Thor_AB_25 and Loki_GC14_75 were tested for the ability to complement the *S. cerevisiae vps4* null mutation. As a control for the Asgard Vps4, we performed the complementation assays with CdvC from *S. solfataricus*_P2 and Bathyarchaeota. Briefly, we re- codon-optimized the coding sequences of Asgard Vps4 and TACK CdvC for expression in *S. cerevisiae*, and, respectively, assembled the coding sequences into transcription units of pPOT-RFP vector that contains a native promoter region of *S. cerevisiae* BY4741 *vps4* (a 500 bp DNA sequence region upstream from the ATG start codon of this gene) and a *S. cerevisiae* Cytochrome c isoform 1 (CYC1) terminator using the YeastFab Assembly method.(31) The assembly products were transformed into *S. cerevisiae vps4Δ* by the LiAc/PEG method(32). As previously described, in *S. cerevisiae, vps4* null mutation resulted in temperature-sensitive growth defect, causing growth arrest at 39 °C (33, 34). We found that the Asgard Vps4 could slightly suppress the growth defect of *vps4Δ* at 39 °C (Fig. 4A). Remarkably, however, after incubation at 39 °C for 96 h, the growth of *vps4Δ* bearing Asgard Vps4 showed substantial, although variable, restoration at 30 °C, in a sharp contrast with *vps4Δ* for which no restoration was observed (Fig. 4A). Nevertheless, both CdvCs showed only minimal growth restoration of *vps4Δ* at 39 °C and a limited enhancement of viability at 30° C; complemetation with these proteins was subtantially less eifficient than that observed with their Asgard counterparts. Furthermore, the *S. cerevisiae* Vps4, Asgard Vps4, and CdvCs were re-codon-optimized, synthesized, and respectively, cloned into a pCold-TF vector (Takara Bio Co Ltd., Japan). After expression in *Escherichia coli* BL21, proteins were purified by Mag-Beads His-Tag Protein Purification Kit (BBI Co., Ltd, China). The biochemical experiments *in vitro* show that these purified proteins are active ATPases both at 30 °C and 39 °C (Fig. 4B). This observation eliminates the possibility that the poor complement result of CdvCs was due to the lack of ATPases activity at 39 °C, and is compatible with the involvement of the ATPase activity of Asgard Vps4 in sustaining the viability of the *S. cerevisiae vps4Δ* mutant under non-permissive conditions.

**FIG 4.**
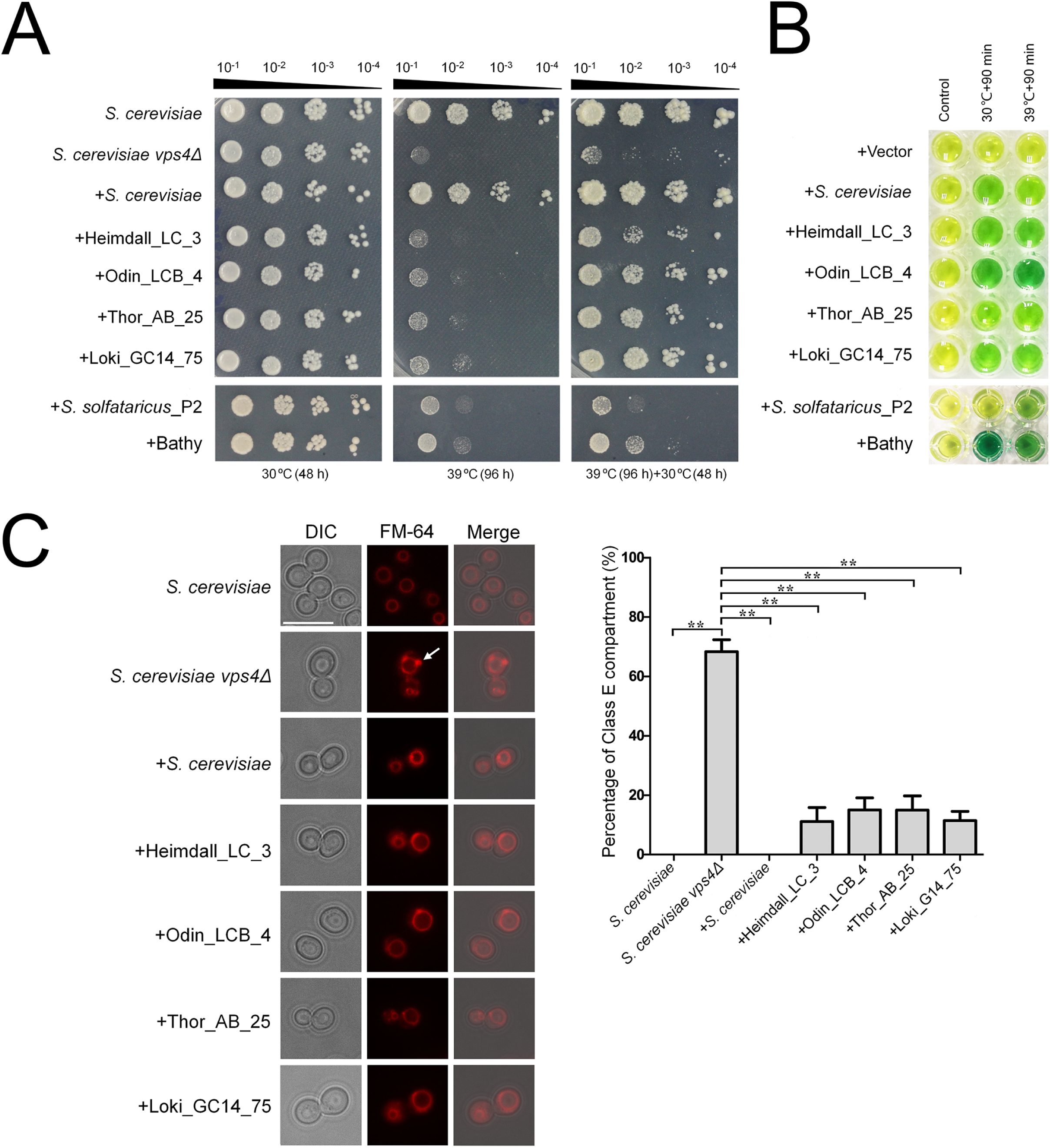
Functional complementation of *Saccharomyces cerevisiae* vps4 null mutants by Asgard Vps4. Complementation of the high-temperature-sensitive growth defect of vps4 mutant cells. Five microliters of a series of 10-fold dilutions derived from a starting suspension of an OD_600_ of 10^−1^ was inoculated into SC-Ura medium. (B) The ATPase activity of *S. cerevisiae* Vps4, Asgard Vps4 and Cdvs at 30 ºC and 39 ºC were, respectively, confirmed by a malachite green assay. The substrates would turn from golden to green owing to the inorganic phosphate relased form ATP hydrolysis by Vps4 under the indicated condition. (C) The class E compartments in *S. cerevisiae* vps4 null mutants were largely abrogated by Asgard Vps4. The vacuolar morphologies in the indicated strains were visualized by fluorescent microscopy. Arrowhead highlights the class E compartment in vps4 null mutant. Scale bar=10 μm. Quantification of class E compartment in the indicated strains. The results represented the means from three independent replicates (20 cells per experiment), and standard deviations are indicated with error bars. Statistical significance was assessed by one-way analysis of variance with Bonferroni’s multiple-comparison test. **, *P*<0.01.

As previously described, *vps4* null mutation could induce an formation of an aberrant prevacuolar compartment adjacent to the vacuoles, known as class E compartment, due to the block of intracellular protein trafficking (3, 30, 33). To further demonstrate that Asgard Vps4 is functionally analogous to its eukaryotic counterpart, we observed the vacuoles in the *S. cerevisiae* cells bearing Asgard Vps4. As expected, the characterized class E compartment vacuolar morphology was clearly observed in the *S. cerevisiae vps4Δ* cells, and this defect was almost completely rescued by *S. cerevisiae* Vps4 (Fig. 4C). We found that the Vps4 of Heimdall_LC_3, Odin_LCB_4, Thor_AB_25, and Loki_GC14_75 also partially complemented the aberrant vacuoles in the *vps4* null mutant, with the reduction of class E compartment of about 80% of that observed with the native *S. cerevisiae* Vps4 (Fig. 4C). However, the enlarged vacuoles induced in *vps4Δ* strain were not markedly eliminated by the Asgard Vps4. Taken together, these findings show that the Asgard Vps4 are functionally more similar to the eukaryotic homologs than homologs from other archaea.

## DISCUSSION

In this work, we combined computational approaches, including sequence comparison, phylogenetic analysis and structural modeling, with genetic and biochemical experiments to investigate the evolutionary and functional relationships between the ESCRT-III machineries of Asgard archaea and eukaryotes. Phylogenetic analyses of both the ESCRT-III subunits and Vps4 ATPase show a clear affinity between Asgard archaea and eukaryotes, to the exclusion of the other archaea. Moreover, the divergence of the two groups of ESCRT-III subunits already occurred in Asgard archaea.

The results of amino acid sequence analysis and structural modeling are best compatible with the coevolution of Vps4 with the ESCRT-III subunits. In particular, the interaction between the MIT domain of Vps4 and the MIM1- and MIM2-like of the ESCRT-III subunits appears to have evolved already in Asgard archaea.

The findings of the computational analysis are complemented by our experimental results. In particular, we show that Asgard Vps4 is capable of complementing the *S. cerevisiae* vps4 null mutant much more efficiently than homologs from Crenarchaeota and Bathyarchaeota. This enhanced functionality might be underpinned by the evolution of distinct, “eukaryote-like” structural features, such as the arginine collar that is involved in the disassembly of ESCRT-III polymers.

Taken together, all these findings are compatible with the direct origin of the eukaryotic ESCRT machinery from the Asgard ancestor. In a broader evolutionary context, the ESCRT complex likely evolved in the common ancestor of the TACK and Asgard superphyla whereas its further elaboration occurred in the Asgard lineage. The key event apparently was the duplication of CdvB that seems to combine features of Vps2/24/46 and Vps20/32/60 (12), with subsequent functional diversification of the subunits and coevolution with Vps4.

An intriguing outstanding question is the function of the ESCRT machinery in the Asgard archaea. There is no indication that these (or any other) archaea possess intracellular membranes (22), so the ECRT-III proteins and Vps4 are likely to be involved in cell division as demonstrated for the Cdv proteins of Crenarchaeota. However, the specialization of the ESCRT-III subunits might provide for the formation of eukaryotic-like filaments that could be involved not only in the inside-out fission to produce membrane vesicles that have been observed in the MK-D1 strain, but also the outside-in fission that allows the Asgard archaea to engulf their bacterial metabolic partners. The latter capability is critical for the ‘Entangle-Engulf- Enslave model’ of eukaryogenesis (22). Further molecular and cell biological study of the Asgard membrane remodeling apparatus, even if challenging due to the recalcitrance of these organisms to growth in culture, should shed light on the origin of the eukaryotic endomembrane system, one of the key aspects of eukaryogenesis.

## MATERIALS and METHODS

### Bioinformatics analysis

All the protein sequences were obtained either by NCBI accession number or by BLAST search(35) the non-redundant protein sequences against local Nr database. The protein sequences were aligned using MUSCLE (V3.8.1551)(36), trimmed with TrimAl (V1.4)(37) before construction of phylogenetic trees using IQ-Tree (V1.6.5)(38). The indicated functional domains of proteins were analyzed by Interpro (https://www.ebi.ac.uk/interpro/) and NCBI’s conserved domain database.

### Homology modeling and docking study

We searched the Vps4, Vps2/24/46, and Vps20/32/60 sequences belonging to *S. cerevisiae*, Lokiarchaeum_GC14_75, Thorarchaeota_AB_25, Heimdallarchaeota_LC_3, and Odinarchaeota_LCB_4 from NCBI database and CdvC sequences belonging to to *Sulfolobus solfataricus*_P2, and Bathyarchaeota (http://www.ncbi.nlm.nih.gov, accession number: KZV07689.1, P36108.2, NP_013794.1; KKK42121.1, KKK42122.1, KKK44605.1; OLS30569.1, OLS30568.1, OLS30800.1; OLS27542.1, OLS27541.1, OLS27540.1; OLS18192.1, OLS18193.1, OLS18194.1, AAK41192.1, WP_119819537.1, respectively) to build homology model. The Cryo-EM structure of Vps4 (E233Q) hexamer belonging to *S. cerevisiae* was obtained from the Protein Data Bank (PDB code: 5XMI)(39), and the subunit B was choosen for modeling template and the missing residues (1-118) were built at the I-TASSER server (http://zhanglab.ccmb.med.umich.edu/I-TASSER). Sequence alignments and homology modelings of Vps4 for Lokiarchaeota, Thorarchaeota, Heimdallarchaeota, and Odinarchaeota with unknown structures were carried out on MODELLER program(40), downloaded and installed from salilab server (https://salilab.org/modeller/download_installation.html). The three-dimensional structures of Vps2/24/46 and Vps20/32/60 for *S. cerevisiae* and four Asgard archaea were also built at the I-TASSER server. Among several three-dimensional models generated using homology modeling and ab initio method, the best model was selected after a series of refining and minimization and molecular dynamic simulation employing ff14SB force fields parameters by AMBER 16.0 package(41). Then the complexes of Vps2/24/46 and Vps20/32/60 against Vps4 were simulated using the ZDOCK server(42). Ten top docking poses were generated.

### Molecular dynamic (MD) simulation

The parallel version of AMBER 16.0 package was used to prepare the complex files and conduct MD simulations employing ff14SB force fields parameters. The ionizable residues default protonation states in AMBER 16.0 were assigned. All MD simulations were carried out by applying cubic periodic boundary conditions (PBC) and in an explicit water box of TIP3P water molecules(43) with a minimum distance of 10.0 Å between complex surface and water box boundary. The Na^+^ or Cl^-^ counterions were added in sufficient number to neutralize any net charges of the structures above. All of chemical bond lengths of hydrogen-heavy atoms were restrained by the SHAKE algorithm(44). A cutoff radius of 10.0 Å was set for both non- bonded electrostatic and van der Waals interactions. Long-range electrostatic forces were taken into account using Particle Mesh Ewald (PME) method(45). Langevin dynamics and Langevin piston methods were applied to keep the temperature (300 K) and the pressure (1 bar) of the system constant, respectively. The time step was set to 2.0 fs.

The solvated systems were minimized using PMEMD.CUDA module enabled NVIDIA graphics processing units (GPUs)(46, 47) in three stage, keeping the solute fixed and just minimized the positions of the water and counterions firstly with 100 kcal/(mol·Å^2^) restraints and then reduced to 10 kcal/(mol·Å^2^), and lastly for the entire system without any restraining force. Each stage was conducted with 10000 steps of steepest descent algorithm followed by 1000 steps conjugate gradient minimization to get rid of any unfavorable steric contacts for both solvent and protein molecules. Then, a NVT simulation was conducted to slowly heat the systems temperature from 0 K to 300 K over a period of 500 ps, and density equilibrated for 2000 ps with a weak restraint applied to the whole protein at 1 atm and 300 K. Finally, all restraints were removed, and production MD simulations were carried out at constant pressure (1 atm) and temperature (300 K) in NPT ensemble. For each system, MD simulation was performed for 500 ns and repeated thrice with different random number, and a total of 1.5 μs trajectory was analyzed by CPPTRAJ module (48).

### Calculations of binding free energy

The binding free energies of Vps2/24/46 and Vps20/32/60 against Vps4 were calculated by molecular mechanics-generalized Born surface area (MM-GBSA) method(28, 29). All energy components were calculated using 500 snapshots that were extracted every 200 ps during the last 100 ns of each MD simulation trajectory. The configurational entropy was not considered in the approach as it is extremely time-consuming. So the binding free energy in the solvent environment can be expressed as:

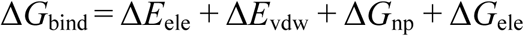

The Δ*E*_ele_, Δ*E*_vdw_, Δ*G*_np_, Δ*G*_ele_ represented electrostatic energy in the gas phase, Van der Waals energy in the gas phase, non-polar solvation energy, and polar solvation energy, respectively. All energy terms were calculated using MM-GBSA calculations, and the Δ*G*_ele_ is estimated by GB model(29), and the Δ*G*_np_ is calculated from the solvent accessible surface area (SASA) of the molecules by molsurf, with the values 0.00542 and 0.92 for SURFTEN and SURFOFF, respectively(49). The decomposition of binding free energies were calculated at residue-pair level for a further investigation of the complexes interactions using the MM-GBSA decomposition program(50, 51) implemented in AMBER 16.0.

### Protein expression in *Escherichia coli* BL21 and purification for biochemical assays *in vitro*

The Vps4, Vps2/24/46, and Vps20/32/60 coding sequences belonging to *S. cerevisiae*, Lokiarchaeum_GC14_75, Thorarchaeota_AB_25, Heimdallarchaeota_LC_3, and Odinarchaeota_LCB_4, and CdvC coding sequences belonging to *Sulfolobus solfataricus*_P2, and Bathyarchaeota from NCBI database (http://www.ncbi.nlm.nih.gov, accession number: KZV07689.1, P36108.2, NP_013794.1; KKK42121.1, KKK42122.1, KKK44605.1; OLS30569.1, OLS30568.1, OLS30800.1; OLS27542.1, OLS27541.1, OLS27540.1; OLS18192.1, OLS18193.1, OLS18194.1, AAK41192.1, WP_119819537.1, respectively) were codon optimized by *GeneDesgin* (http://54.235.254.95/gd/) for expression in *E. coli* BL21, synthesized (BGI Genomics Co., Ltd), and, respectively, cloned into a pCold-TF vector (Takara Bio Co Ltd., Japan) that includes an N-terminal His tag and a soluble trigger factor chaperone tag. The *E. coli* BL21 (Takara Bio Co Ltd, Japan) beared the recombinant vectors were inoculated in LB medium containing 100 μg/ml carbenicillin, and incubated at 37°C until the OD_600_ reached at 0.6-0.8, and then the isopropyl-d-1-thiogalactopyranodside was added at the final concentration of 0.5 mM, followed by incubation at 15 °C for 18-24 h. The cells pellets were collected and resuspended in 20 ml binding buffer (20 mM phosphate buffer (pH 7.4), 500 mM NaCl, 50 mM imidazole, 1 mM dithiothreitol, 1 mM lysozyme, and 1 mM phenylmethylsulfonyl fluoride), followed by ultrasonic decomposition. Next, the target proteins were purified by Mag-Beads His-Tag Protein Purification Kit (BBI Co., Ltd, China) with wash buffer (20 mM phosphate buffer (pH 7.4), 500 mM NaCl, 100 mM imidazole, and 0.1% NP-40) and elution buffer (20 mM phosphate buffer (pH 7.4), 500 mM NaCl, and 500 mM imidazole). Finally, the purified proteins were concentrated to 1-2 ml in phosphate buffered saline (PBS, pH 7.4) by 30K Amicon Ultra-15 (Millipore Co Ltd., USA). Concentrations of these proteins were determined by Bradford Protein Assay Kit (Beytotime Bio Co Ltd., China). The purified Vps4, Vps2/24/46, and Vps20/32/60 belonging to *S. cerevisiae*, Lokiarchaeum_GC14_75, Thorarchaeota_AB_25, Heimdallarchaeota_LC_3, and Odinarchaeota_LCB_4 were used for Isothermal titration calorimetry assay. The purified Vps4 belonging to *S. cerevisiae*, Lokiarchaeum_GC14_75, Thorarchaeota_AB_25, Heimdallarchaeota_LC_3, and Odinarchaeota_LCB_4, and Cdv belonging to *Sulfolobus solfataricus*_P2, and Bathyarchaeota were used for ATPase activity assay.

### Isothermal titration calorimetry assay

Isothermal titration calorimetry assay (ITC) was carried out at 25°C using an ITC200 system (MicroCal, USA). The Vps4 MIT domain (3 μM in PBS buffer) was placed in cell and titrated with 19 injections of 10 μl of Vps2/24/46 or Vps20/32/60 (33 μM in PBS buffer) at 2 min intervals. The heat of ligand dilution into buffer was subtracted from the reaction heat, after removing the data of 1^st^ injection. Data analysis was carried out using Origin 7.0 (MicroCal, USA).

### ATPase activity assay

The ATPase activity was determined by a slightly modified malachite green assay(52). In short, the purified proteins (4 μM) were incubated with reaction buffer (1 mM ATP, 20 mM HEPES pH 7.4, 100 mM NaCl, 10 mM MgCl_2_, 1 mM DTT) in a total volume of 50 μl at the indicated temperature for 90 min, and was immediately stopped by liquid nitrogen. Then, the reaction mixture was added with 100 μl of malachite green color buffer (14 mM ammonium molybdate, 1.3 M HCl, and 1.5 mM malachite green) and 50 μl of 21% (w/v) citric acid, followed by incubation at room temperature for 30 min. Finally, the reaction mixture that turned green was attributed to the free phosphate released by Vps4 ATP hydrolysis. Additionally, the control experiments were identical to the treatment group, except that the mixture of Vps4 and reaction buffer was immediately treated with liquid nitroge before addition of malachite green color buffer and citric acid; and these experiments were to eliminate the interference of irrelevant free phosphate. Also, the empty vector was served to prove that the ATP hydrolysis is due to Vps4.

### *S. cerevisiae* strains and cultivation

The *S. cerevisiae* strain BY4741 (*MATa leu2Δ0 met15Δ0 ura3Δ0 his3Δ1*) and its derivative *vps4* null mutant strain YPR173Ca (designated the *S. cerevisiae*Δ*vps4* in this study) were from *S. cerevisiae* deletion mutant library (53). *S. cerevisiae* cells were routinely cultured in YPD medium (10 g/L yeast extract, 20 g/L peptone, 20 g/L glucose) or SC-Ura medium (6.7 g/L YNB, 0.01μmol/L Fe(NH_4_)_2_(SO_4_)_2_, 20 g/L glucose, and complete amino acids without uracil) at 30 ºC unless otherwise noted. The solid media were identical to those of YPD or SC-Ura except that agar was present.

### Complementation Assay

The Vps4 coding sequences belonging to Lokiarchaeum_GC14_75, Thorarchaeota_AB_25, Heimdallarchaeota_LC_3, and Odinarchaeota_LCB_4, and CdvC coding sequences belonging to *Sulfolobus solfataricus*_P2, and Bathyarchaeota (NCBI accession number: KKK42121.1, OLS30569.1, OLS27542.1, OLS18192.1, AAK41192.1, WP_119819537.1, respectively) were codon optimized by *GeneDesgin* (http://54.235.254.95/gd/) for expression in *S. cerevisiae*, before synthesis by BGI Genomics Co., Ltd (54). To eliminate the interference of transcriptional level factor, a native promoter region of *S. cerevisiae* BY4741 *vps4* (a 500 bp DNA sequence region upstream from the ATG start codon of this gene) was used to drive the coding sequences. Then, we assembled the coding sequences, the *S. cerevisiae vps4* native promoter, and a *S. cerevisiae* CYC1 (Cytochrome c isoform 1) terminator into a pPOT-RFP vector according to a developed YeastFab Assembly protocol(31). Besides, the pPOT-RFP vector containing the entire *S. cerevisiae* BY4741 *vps4* with its native promoter and the CYC1 terminator were transformed into the *vps4* null mutant *S. cerevisiae*(32), and this reconstituted strain was designated the “+*S. cerevisiae*". In this study, both the *S. cerevisiae* and *S. cerevisiae vps4Δ* were transformed with the pPOT-RFP vector as the control.

### FM-64M staining

*S. cerevisiae* cells of each strain were cultured in SC-Ura medium at 30 ºC and normalized to an OD_600_ of 0.5-0.8. Then, the *S. cerevisiae* cells were stained with 80 μM FM-64M (AAT Bioquest Co Ltd., China) at 30 ºC for 20 min, and next cultured for 120 min after washes with medium. Finally, the *S. cerevisiae* cells were examined under a N-STORM fluorescence microscope (Nikon Co Ltd., Japan).

## ACKNOWLEDGMENT

This work was supported by the National Natural Science Foundation of China (Grant Nos. 91851105, 31622002, 31970105, 31725002, 21625302, 31800615), the China Postdoctoral Science Foundation (Grant No. 2018M643153), the Basic and Applied Basic Research of Guangdong Province (Grant No. 2019A1515110089), the Shenzhen Science and Technology Program (Grant No. JCYJ20170818091727570, KQTD20180412181334790), Shenzhen Key Laboratory of Synthetic Genomics (ZDSYS201802061806209), Guangdong Provincial Key Laboratory of Synthetic Genomics (2019B030301006) and the Key Project of Department of Education of Guangdong Province (No. 2017KZDXM071). EVK is supported by the Intramural Research Program funds of the National Institutes of Health of the USA.

## AUTHOR’ CONTRIBUTIONS

Zhongyi Lu and Meng Li conceived and designed the experiments. Zhongyi Lu, Tianyi Li, Siyu Zhang, and Jinquan Li performed the experiments. Huiying Chu and Guohui Li designed the molecular dynamics strategy. Ting Fu performed the simulations. Zhongyi Lu, Ting Fu, and Huiying Chu analyzed the data. Yang Liu and Junbiao Dai contributed reagents/materials/analysis tools. Zhongyi Lu, Ting Fu, Eugene Koonin and Meng Li wrote and all authors edited and approved the paper.

## Supplemental information

**FIG S1. Phylogenetic analysis of ESCRT-III-related proteins in the eukarya and archaea**. The tree was reconstructed by maximum likelihood analysis using 156 representative amino acid sequences based on LG+G4 model (recommended by the “TESTONLY”), with option “-bb 1000", and the bootstrap values are shown on nodes.

**FIG S2. Phylogenetic analysis of Vps4-related proteins in eukarya and archaea**. The tree was reconstructed by maximum likelihood analysis using 76 representative amino acid sequences based on LG+I+G4 model (recommended by the “TESTONLY”), with option “-bb 1000", and the bootstrap values are shown on nodes.

**FIG S3. Predicted** “**arginine collar**” **in Vps4 of Asgard archaea and eukarya**. (A) The Walker A, Walker B, Sensor I, ARG finger, and Sensor II are conserved across all the indiactaed sequences, and are shown to confirm the location of “arginine collar". Conserved arginine residues of “arginine collar” are highlighted (red shading and red letters, respectively). The information of proteins used here can be found in Table S1. (B) The top and bottom views of the hexameric ring (grey) were constructed by Heimdall_LC_3 Vps4 (white) as the example to demonstrate the location of “arginine collar", including R222, R231, and R232.

**FIG S4. Phylogenetic analysis of the Microtubule Interacting and Transport domain in Vps4-related proteins**. The tree was reconstructed by maximum likelihood analysis using 48 amino acid sequences based on LG+I+G4 model (recommended by the “TESTONLY”), with option “-bb 1000", and the bootstrap values are shown on nodes.

**FIG S5. The number of Vps4 and CdvC Microtubule Interacting and Transport domain amino acid residues in alpha conformation of Asgard archaea, *Saccharomyces cerevisiae* and TACK archaea during molecular dynamic simulations**. The curves of the numbers of Vps4 MIT domain amino acid residues in alpha conformation of Asgard and *S. cerevisiae*, which were calculated from the last 200 ns MD simulation trajectories, were plotted against simulation time.

**FIG S6. ITC binding profiles of Asgard Vps4 Microtubule Interacting and Transport domain titrated with Asgard Vps2/24/46 and Vps20/32/60**. (A) The curve of Heimdall_LC_3 TF-Vps4-MIT titrated with Heimdall_LC_3 TF-Vps2/24/46 was fit to Sequential Binding Sites; ΔH_1_=- 4.73×10^5^ cal mol^-1^; ΔH_2_=9.31×10^5^ cal mol^-1^. The curve of Heimdall_LC_3- Vps4-MIT titrated with Heimdall_LC_3 TF-Vps20/32/60 was fit to Sequential Binding Sites; ΔH_1_=2.78×10^6^ cal mol^-1^; ΔH_2_=-2.31×10^6^ cal mol^-1^. (B) The curve of Odin_LCB_4 TF-Vps4-MIT titrated with Odin_LCB_4 Vps2/24/46 was fit to One Set of Sites; ΔH=3.92×10^5^ cal mol^-1^. The curve of Odin_LCB_4 TF-Vps4-MIT titrated with Odin_LCB_4 TF-Vps20/32/60 was fit to One Set of Sites; ΔH=3.31×10^5^ cal mol^-1^. (C) The curve of Thor_AB_25 TF-Vps4-MIT titrated with Thor_AB_25 TF-Vps2/24/46 was fit Sequential Binding Sites; ΔH_1_=3.25×10^8^ cal mol^-1^; ΔH_2_=1.29×10^6^ cal mol^-1^. The curve of Thor_AB_25 TF-Vps4-MIT titrated with Thor_AB_25 TF-Vps20/32/60 was fit to One Set of Sites; ΔH=1.62×10^6^ cal mol^-1^. (D) The curve of Loki_GC14_75 TF-Vps4-MIT titrated with Loki_GC14_75 TF-Vps2/24/46 was fit to Sequential Binding Sites; ΔH_1_=-1.21×10^5^ cal mol^-1^; ΔH_2_=4.22×10^5^ cal mol^-1^. The curve of Loki_GC14_75 TF-Vps4-MIT titrated with Loki_GC14_75 TF-Vps20/32/60 was fit to One Set of Sites; ΔH=1.42×10^6^ cal mol^-1^. Binding to a TF control surface was negligible (not shown).

**TABLE S1. Summary of proteins used in this study**.

**TABLE S2. The predicted binding free energies between Vps4 and ESCRT-III subunits (Vps2/24/46 (A), and Vps20/32/60 (B))**.

**TABLE S3. The dominant amino acid residues of Vps4 Microtubule Interacting and Transport domain involved in binding with Vps2/24/46 are listed**.

**TABLE S4. The dominant amino acid residues of Vps4 Microtubule Interacting and Transport domain involved in binding with Vps20/32/60 are listed**.

